# Empowering Multi-Cohort Gene Expression Analysis to Increase Reproducibility

**DOI:** 10.1101/071514

**Authors:** Winston A. Haynes, Francesco Vallania, Charles Liu, Erika Bongen, Aurelie Tomczak, Marta Andres-Terrè, Shane Lofgren, Andrew Tam, Cole A. Deisseroth, Matthew D. Li, Timothy E. Sweeney, Purvesh Khatri

## Abstract

A major contributor to the scientific reproducibility crisis has been that the results from homogeneous, single-center studies do not generalize to heterogeneous, real world populations. Multi-cohort gene expression analysis has helped to increase reproducibility by aggregating data from diverse populations into a single analysis. To make the multi-cohort analysis process more feasible, we have assembled an analysis pipeline which implements rigorously studied meta-analysis best practices. We have compiled and made publicly available the results of our own multi-cohort gene expression analysis of 103 diseases, spanning 615 studies and 36,915 samples, through a novel and interactive web application. As a result, we have made both the process of and the results from multi-cohort gene expression analysis more approachable for non-technical users.

## 1. Introduction

Prior to translation of the results of a biological experiment into clinical practice, they must be replicated and validated in multiple independent cohorts. However, the majority of findings fail to validate, leading to a ‘reproducibility crisis’ in science.^1,2^ One of the factors in this lack of reproducibility is that traditional, single cohort studies do not represent the heterogeneity observed in the real world patient population.^3^ As a result, observed and reported effects are often specific to a population subset instead of generalizable across the population.

More than two million publicly available gene expression microarrays present novel opportunities to incorporate the real-world heterogeneity observed in patient populations into analysis.^4,5^ However, the biological (tissue, treatment, demographics) and technical (experimental protocol, microarray) heterogeneity present in such data poses a daunting challenge for their integration and reuse. Consequently, many tools, which allow reuse of these data, are unable to combine evidence across multiple data sets and place that burden on the end user, leading to under-utilization of these datasets.^6,7^

Previously, we have described a novel multi-cohort analysis framework for integrating multiple heterogeneous datasets to identify robust and reproducible signatures by leveraging the biological and technical heterogeneity in these datasets. We have repeatedly demonstrated the utility of our framework for identifying novel diagnostic and prognostic biomarkers, drug targets, and repurposing FDA-approved drugs in diverse diseases, including organ transplantation, cancer, infection, and neurodegenerative diseases.^8–16^ In each of these analyses, we analyzed more than a thousand human samples from more than 10 independent cohorts to generate and validate data-driven hypotheses. Many of these results also been further validated in experimental settings.^8,11,16^ These results have further demonstrated the ability of our framework to create “Big Data” by combining multiple smaller studies that are collectively representative of the real word patient population heterogeneity.

## 2. Multi-Cohort Gene Expression Analysis with MetaIntegrator

Despite its demonstrated utility in identifying robust, reproducible, and biologically as well as clinically relevant disease signatures, our multi-cohort analysis framework has previously required manual dataset download, pipeline set up, and visualization generation. To lower this barrier to entry, we have developed MetaIntegrator, an R package that automates most of the multi-cohort analysis framework. Our package guides the user from data download to execution of statistical analysis to evaluation of the results [Figure 1].

**Fig. 1.**
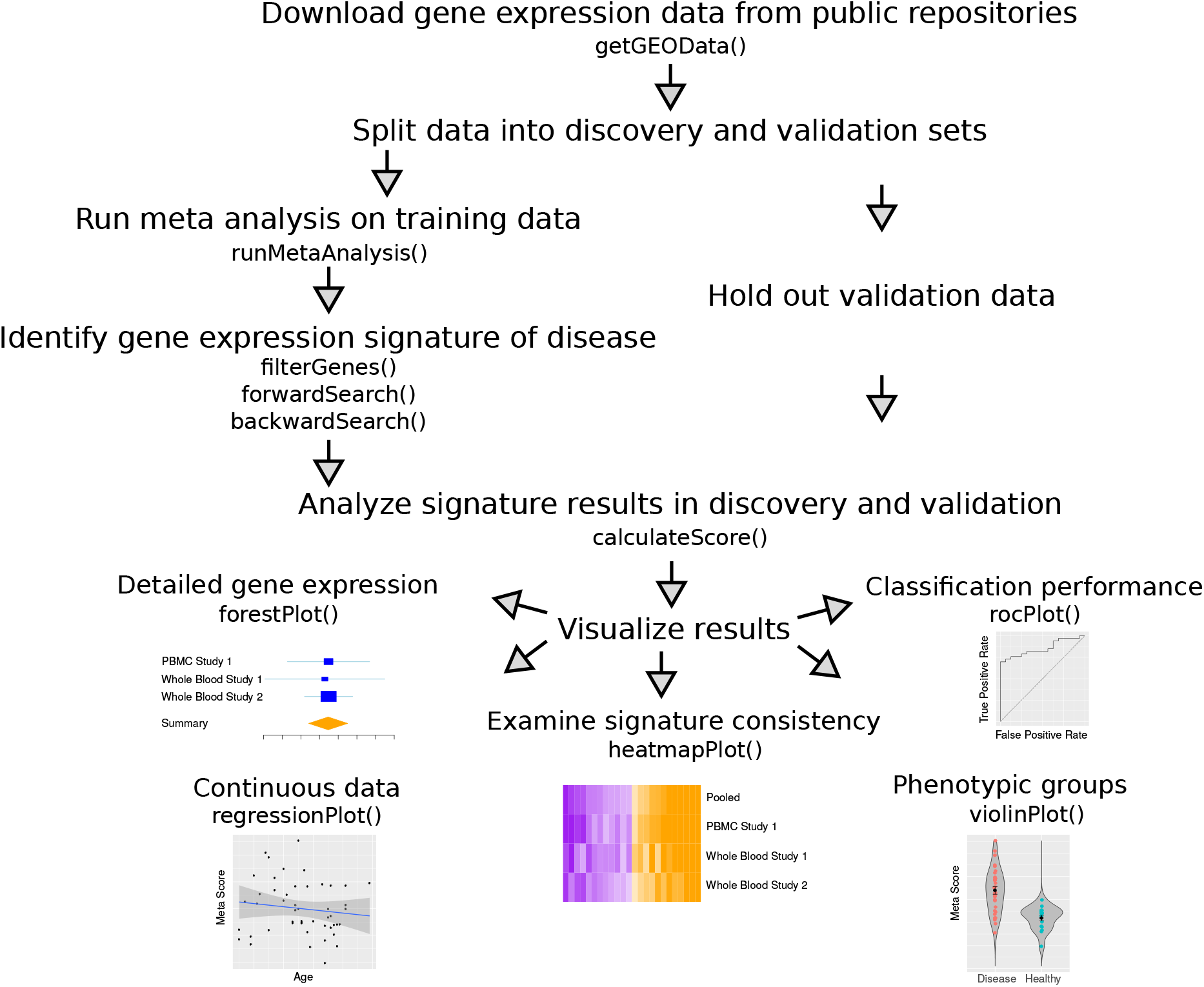
Gene expression meta-analysis workow with MetaIntegrator utility functions.

### 2.1. Data Processing

The first step in the multi-cohort analysis is downloading the requisite experimental information, notably the class labels (case or control), the gene expression data, and any interesting phenotypic information about the samples. Since we have found that most users will download data from the NCBI’s Gene Expression Omnibus (GEO), we have integrated an automatic downloading and processing of GEO data into our analysis pipeline. MetaIntegrator will automatically download the expression data and all available annotations, perform sanity checks that the data have been appropriately normalized, and compile the data into the MetaIntegrator object format.

### 2.2. Multi-cohort Analysis

#### 2.2.1. Combining effect sizes

Our meta-analysis approach computes an Hedges g effect size for each gene in each dataset defined as:

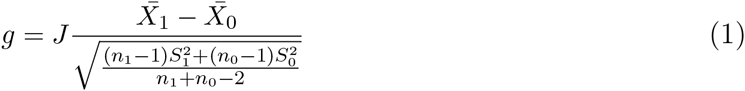

where 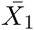 and 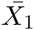 are the average expression for cases and controls, *S*_1_ and *S*_0_ are the standard deviations for cases and controls, and *n*_1_ and *n*_0_ are the number of cases and controls.^8,17^ *J* is the Hedges’ g correction factor, which is computed as:

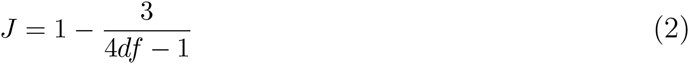

where *df* are the degrees of freedom.

To pool these effect sizes across datasets, the summary effect size *g*_*s*_ is computed using a random effect model as:

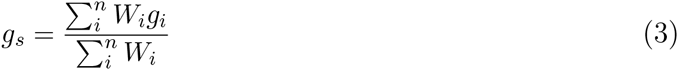

where *n* is the number of studies, *W*_*i*_ is a weight equal to 1/(*V*_*i*_+*T*^2^), *V*_*i*_ is the variance of that gene within a given dataset *i*, and *T*^2^ is the inter-dataset variation as estimated by the DerSimonian-Laird method.^17,18^ The standard error for the summary effect size is 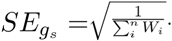. Given *g*_*s*_ and *SE*_*gs*_, we calculate a p-value based on a standard normal distribution and perform a Benjamini-Hochberg FDR correction for multiple hypothesis testing.^19^

#### 2.2.2. Heterogeneity of effect size

We calculate Cochrane’s Q value for evaluating heterogeneity of effect size estimates between studies:

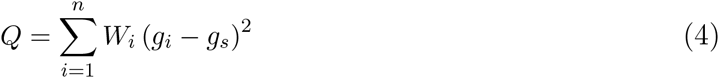

where *W*_*i*_, *g*_*i*_, and *g*_*s*_ are the same as above.^17^ The p-value of Cochrane’s Q is calculated against a chi-squared distribution and adjusted for multiple hypothesis testing using the Benjamini-Hochberg FDR method.^19^ A statistically significant Cochrane’s Q indicates significant heterogeneity of effect sizes between studies.

#### 2.2.3. Combining p-values

We use Fisher’s method for combining p-values across studies.^20^ We calculate the log sum of p-values that each gene is up-regulated as:

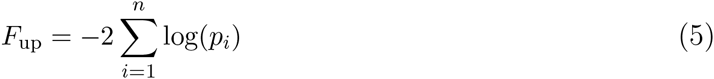

where *n* is the number of studies and *p*_*i*_ is the t-test p-value that the gene of interest is up-regulated in study *i*. Similarly, we calculate *F*_down_ as the log-sum of p-values that each gene is down-regulated.

For each gene, we calculate the p-value of *F*_up_ and *F*_down_ under a chi-squared distribution and perform a Benjamini-Hochberg FDR correction.^19^

### 2.3. Signature Selection

Once meta-analysis is performed, a subset of genes must be identified as the disease signature. MetaIntegrator allows the user to identify these genes by varying the filtering parameters based on gene effect size, effect size false discovery rate, Fisher’s method false discovery rate, heterogeneity of effect size, and the number of studies in which the gene was present. In order to avoid disproportionate influence of a single study, MetaIntegrator allows the user only include genes which were similarly significant across all leave-one-dataset-out analyses. By varying these criterion, the user may control whether they identify a larger set of genes, which may be ideal for understanding molecular pathogenesis and identifying drug targets, or a smaller set of genes, which may be optimal developing a parsimonious clinical diagnostic.

For users that are particularly interested in developing a powerful diagnostic, we have integrated forward and backward search, which reduce gene set size to optimize the area under the receiver operating characteristic curve on the training data.^10^

### 2.4. Score Calculation

For a set of signature genes, a signature score can be computed for every sample, *i*, as:

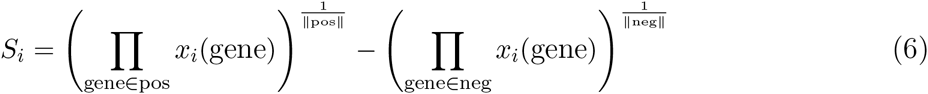

where pos and neg are the sets of positive and negative genes, respectively, and *x*_*i*_(gene) is the expression of any particular gene in sample *i* (a positive score indicates an association with cases and a negative score with controls). This score *S*_*i*_ is normalized to a z-score to center the samples for each study around zero.

### 2.5. Visualization

With scores calculated for each sample, we are able to visualize comparisons of cases vs. controls, regression of continuous variables against the score, and consistency of gene expression across datasets. Some of the built in visualizations, in counter-clockwise order from Figure 1:

- **Forest plots.** Examine the effect sizes and standard errors for a single gene across studies, including the summary effect size.
- **Regression plots.** Evaluate the relationship of the signature score with continuous variables like clinical severity and time.
- **Heatmap plots.** Observe consistency of differential expression for all signature genes across studies.
- **Violin plots.** Compare signature scores across categorical variables like disease sub-types, treatment protocols, and demographic groups.
- **ROC plots.** Evaluate classification performance for signature score on a single dataset in terms of specificity and sensitivity.

## 3. Data-Driven Biological Hypotheses with MetaSignature

We have created MetaSignature (**http://metasignature.stanford.edu**), a web application which empowers researchers to generate data-driven hypotheses by enabling access to the results of our multi-cohort gene expression analysis framework. We focused on enabling intuitive data access for researchers with specific interest in either a disease, a gene, or several genes, while requiring little or no analytic background.

### 3.1. Data

Thus far, we have aggregated 615 gene expression studies composed of more than 35,000 human samples with approximately 1.5 billion data points from 103 diseases, a number which we will continue to grow. For each disease, we applied our multi-cohort analysis approach to compute the gene expression differences between the manually curated cases and controls. To perform these multi-cohort analyses, we searched for relevant studies in GEO, identified cases and controls in every study, and calculated disease effect sizes using the MetaIntegrator R package. We stored the multi-cohort analysis results in a MySQL database for rapid retrieval. As more studies are incorporated into our database, we recalculate the disease summary effect sizes.

### 3.2. Gene-centric Analysis

For researchers that are interested in the expression of a particular gene, MetaSignature provides visualizations that allow researchers to quickly identify the diseases in which specified gene is most differentially expressed [Figure 2a], study-level data of the gene expression in particular diseases [Figure 2b], and cell type-specific gene expression patterns [Figure 2c].

**Fig. 2.**
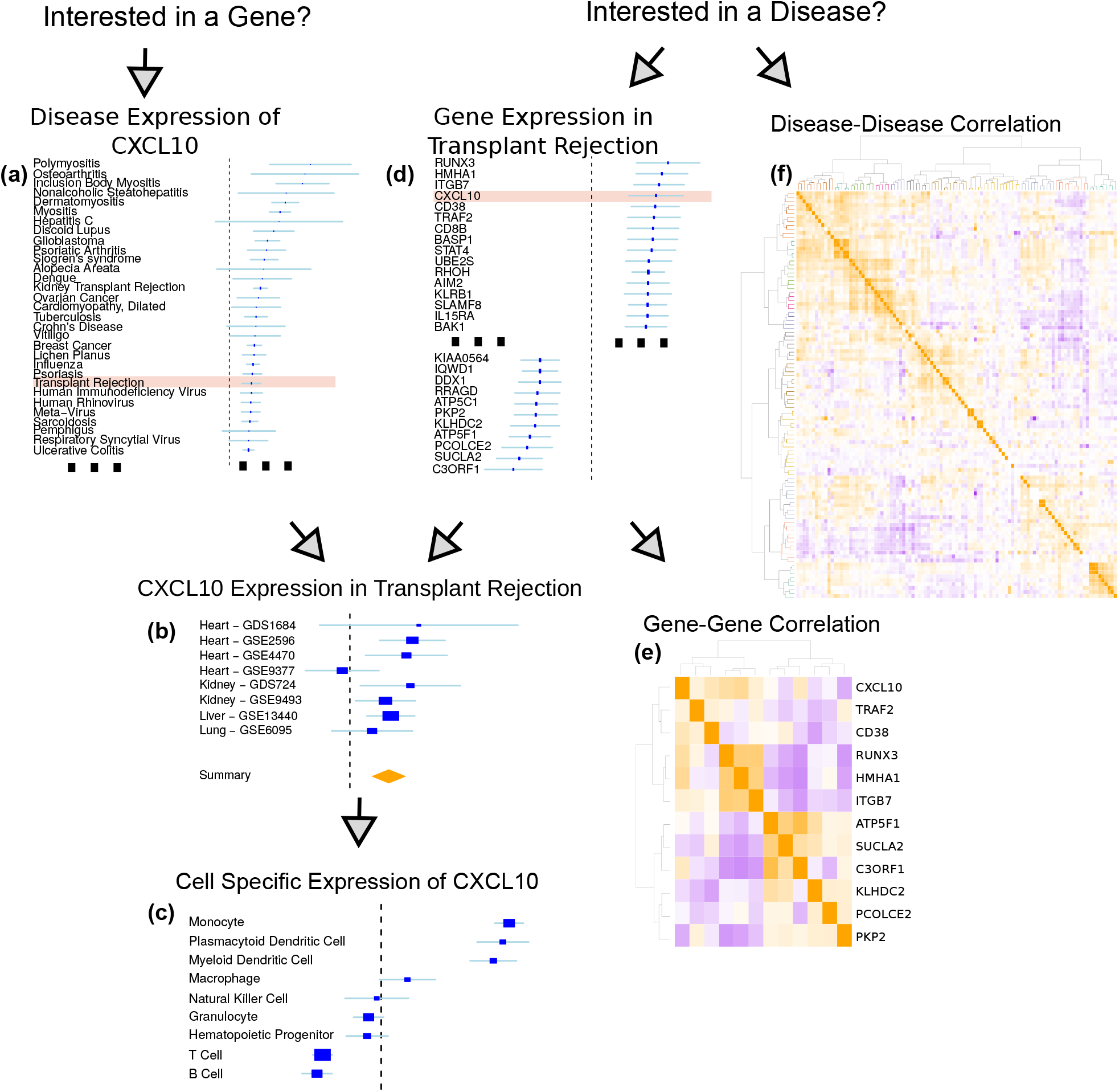
Diagram of the MetaSignature web application.

For instance, consider a researcher who has developed a drug, such as atorvastatin, that effectively reduces plasma levels of *CXCL10*, and seeks to identify the most promising clinical applications. Using MetaSignature, she determines *CXCL10* is significantly up-regulated in transplant rejection [Figure 2a]. A drilldown further identifies eight separate studies that have measured *CXCL10* in transplant rejection, indicating a highly positive effect size in all except one of these studies [Figure 2b]. The researcher further observes that *CXCL10* is up-regulated in monocytes, compared to other immune cell types. [Figure 2c]. Taken together, these findings would motivate a clinical investigation of the use of a *CXCL10* inhibitor, such as atorvastatin, in monocytes of patients at risk for transplant rejection. We have already verified this data-driven hypothesis in mouse models and patient electronic health records, where, in both cases, atorvastatin increases graft survival.^8^

Beyond single gene analysis, MetaSignature empowers users to examine gene sets in terms of correlation of those genes based on their disease effect sizes [Figure 2e] and correlation of diseases based on expression of that set of genes [similar to Figure 2f]. These visualizations enable dissection of positively- and negatively-correlated members of gene families.

### 3.3. Disease-centric Analysis

If a researcher is more interested in a particular disease, then MetaSignature enables identification of genes that are most up- or down-regulated in that disease [Figure 2d] and exploration of that disease’s relationship to other diseases based on gene expression [Figure 2f]. When we compute disease-disease correlation based on gene expression data, we observe clustering patterns that map to established disease categories.

To follow our example from the gene-centric analysis, consider a researcher who is interested in improving transplant rejection outcomes. To gain a global understanding of transplant rejection, the researcher observes that transplant rejection falls into a cluster of inflammatory diseases, including discoid lupus, Crohn’s disease, and interstitial cystitis [Figure 2f]. By examining the transplant rejection expression data in MetaSignature, he would recognize that *CXCL10*, a chemokine important in inflammatory response, is one of the most up-regulated genes in transplant rejection [Figure 2d].^21^ After verifying that this observation is consistent across studies [Figure 2b], the researcher identifies that *CXCL10* is a reasonable target for therapeutic inhibition in transplant rejection. Looking at other genes which are up- and down-regulated in transplant rejection, he recognizes that *CXCL10* expression is in a positively correlated with several other genes, including *TRAF2* and *CD38* [Figure 2e]. Collectively from these observations, the researcher has learned that transplant rejection is related to inflammatory diseases, which is consistent with the observed up-regulation of CXCL10, an inflammatory chemokine. As noted in the gene-centric analysis above, we have observed increased graft survival through administration of atorvastatin.^8^

## 4. Discussion

The reproducibility crisis in biomedical research has led to erroneous conclusions and wasted resources. Here, we present a vertically integrated platform that can both assist with gene expression multi-cohort analysis (MetaIntegrator) and provide aggregated results for users who wish to rapidly test hypotheses (MetaSignature). By leveraging the growing public data available for study, this new resource can drastically reduce the time and effort for biological hypothesis testing across numerous studies and diseases. While many software packages exist for similar analyses,^22–26^ ours offers simple, custom software for plotting and analysis, automated downloading of data from GEO, and integration to the MetaSignature database.

Our package is complementary to the recently published OMiCC platform, which enables curation and meta-analysis of GEO studies.^27^ OMiCC relies on the RankProd package for performing meta-analysis using rank-based statistics for identifying differentially expressed genes.^28^ While others have provided thorough comparisons of the different meta-analysis methods, the most notable difference between RankProd and MetaIntegrator is that rank-based statistics fail to produce a summary effect size across multiple studies.^29,30^ By leveraging our MetaIntegrator package, OMiCC could produce differential gene expression profiles across multiple studies instead of internal to single studies.

Our work promises to increase reproducibility of research for both data analysts and wet lab researchers. For data analysts, we have made multi-cohort gene expression analysis publicly available through a straightforward R package. By performing integrative, multi-cohort analyses, these analysts will generate more reproducible results. For wet lab researchers, we are empowering data-driven hypotheses prior to experimentation. Rather than performing broad assays to identify disease related genes, researchers can focus on performing targeted experiments on genes which are reproducible across cohorts.

## 5. Package and Source Code Distribution

The MetaIntegrator R package, including an introductory vignette, may be installed using the following command in R:

`install.packages(“MetaIntegrator”)`

The source code for MetaIntegrator is available at: https://cran.rstudio.com/web/packages/MetaIntegrator/

MetaSignature was developed using R and Shiny and is hosted at: http://metasignature.stanford.edu/

